# Potassium starvation induces autophagy in yeast

**DOI:** 10.1101/2020.06.30.179085

**Authors:** Nambirajan Rangarajan, Ishani Kapoor, Shuang Li, Peter Drossopoulos, Kristen K. White, Victoria J. Madden, Henrik G. Dohlman

**Author notes:** Undergraduate researcher (Ishani Kapoor, Peter Drossopoulos). **To whom correspondence should be addressed:** Henrik G. Dohlman, University of North Carolina at Chapel Hill, 4016 Genetic Medicine Building, 120 Mason Farm Rd., Chapel Hill, NC 27599 USA.

## Abstract

Autophagy is a conserved process that recycles cellular contents to promote survival. Although nitrogen starvation is the canonical inducer of autophagy, recent studies have revealed several other nutrients important to this process. In this study, we used a quantitative, high-throughput assay to identify potassium starvation as a new and potent inducer of autophagy. We found that potassium-dependent autophagy requires the core pathway kinases Atg1, Atg5, Vps34, as well as other components of Phosphatidylinositol 3-kinase Complex I. Transmission electron microscopy revealed abundant autophagosome formation in response to both stimuli. RNA sequencing indicated distinct transcriptional responses – nitrogen affects transport of ions such as copper while potassium targets the organization of other cellular components. Thus, nitrogen and potassium share the ability to influence metabolic supply and demand but do so in different ways. Both inputs promote catabolism through bulk autophagy, but inhibit cellular anabolism through distinct mechanisms.

## INTRODUCTION

Many eukaryotic organisms experience nutrient starvation and their ability to adapt is important for survival. Adaptation to starvation is often characterized by alterations to signaling, transcription and metabolism (1-4). To support these changes, cellular components are recycled into useable building blocks by two distinct and complementary mechanisms. Whereas the ubiquitin proteasome system breaks down specific short-lived proteins into their constituent amino acids, autophagy targets a wider variety of cytoplasmic cargo for degradation (5).

Much of our understanding of proteostasis comes from genetic studies conducted in the budding yeast *Saccharomyces cerevisiae*. As first shown in yeast, and later in animals, autophagy can be induced by diverse nutritional and pharmacological signals that converge at the target of rapamycin complex 1 (TORC1) (6-10). In nutrient-rich environments, TORC1 remains active and inhibits autophagy via phosphorylation. Under pro-autophagy conditions, deactivation of TORC1 promotes assembly of autophagy-related (ATG) proteins (such as Atg1) and lipid chains. This step requires the phosphatidylinositol 3-kinase (PI 3-kinase) Vps34, in complex with Vps15 (regulatory kinase), Vps30 (adaptor) and either Atg14 or Vps38 (11-18). Activation of Vps34 leads to increased levels of Phosphatidylinositol 3-phosphate, which enables downstream proteins such as Atg5 to assemble into a functional complex and catalyze the conjugation of the ubiquitin-like protein Atg8 (LC3, GABARAP and GATE-16 in animal cells) to phosphatidylethanolamine (19). This conjugate leads to membrane expansion around portions of the cytoplasm (20). This structure, known as the autophagosome, is a unique double-membrane vesicle that fuses with the vacuole (lysosome, in higher eukaryotes), resulting in the degradation and reuse of cytoplasmic contents (21-26).

In addition to removing cytoplasmic proteins, autophagy helps to replenish the cellular pool of biologically important metals, which are required in large abundance (calcium, potassium, sodium) or in trace amounts (iron, copper, zinc) to maintain physiological parameters such as cell volume, pH and protein synthesis (27-29). Under basal conditions, iron is recycled through the vacuolar transport and degradation of iron-containing cargo (30). Nitrogen starvation results in the release of calcium ions from lysosomes and the subsequent induction of autophagy through the activation of the transcription factor EB (31,32). In addition, the availability of certain ions influences basal as well as stress-induced autophagy. For example, zinc starvation promotes autophagic targeting of zinc-binding proteins in yeast (33). Therefore, autophagy and ion homeostasis share a complex reciprocal relationship that is only beginning to be appreciated.

Here we describe a new and important regulator of autophagy. Using a panel of independent and complementary methods, we show that short-term potassium starvation induces bulk autophagy. Using RNA sequencing, we demonstrate that nitrogen and potassium deprivation result in substantially different transcriptional profiles. While nitrogen affects genes important to ion transport, potassium is less specific and regulates genes related to cellular organization. Taken together, our findings point to converging mechanisms for autophagic recycling of cellular materials in conjunction with distinct and complementary routes to transcriptional adaptation.

## RESULTS

### Potassium starvation promotes autophagy

Nitrogen limitation has long been the canonical inducer of autophagy. However, recent reports suggest other salts, amino acids and micronutrients are also important (33-36). Autophagy may indeed be upregulated to compensate for limited external availability of essential nutrients. We hypothesized that a quantitative and parallel analysis of growth medium components might reveal new pro-autophagy regulatory pathways.

To enable high-throughput comparison of autophagy responses across diverse nutritional conditions, we used Rosella, a fluorescent reporter of autophagy (**Figure 1A**) (37,38). Rosella is comprised of super-ecliptic pHluorin (green, pH sensitive) fused with DsRed.T3 (red, pH stable). Upon induction of autophagy, the reporter is transported to the lumen of the vacuole where the acidic pH lowers green fluorescence, while red fluorescence remains unaffected. Rosella response is presented as a ratio of red and green fluorescence. As shown in **Figure 1B**, cells transferred to low nitrogen conditions exhibited a large increase in Rosella response, as compared to control cells in standard medium.

**Figure 1:**
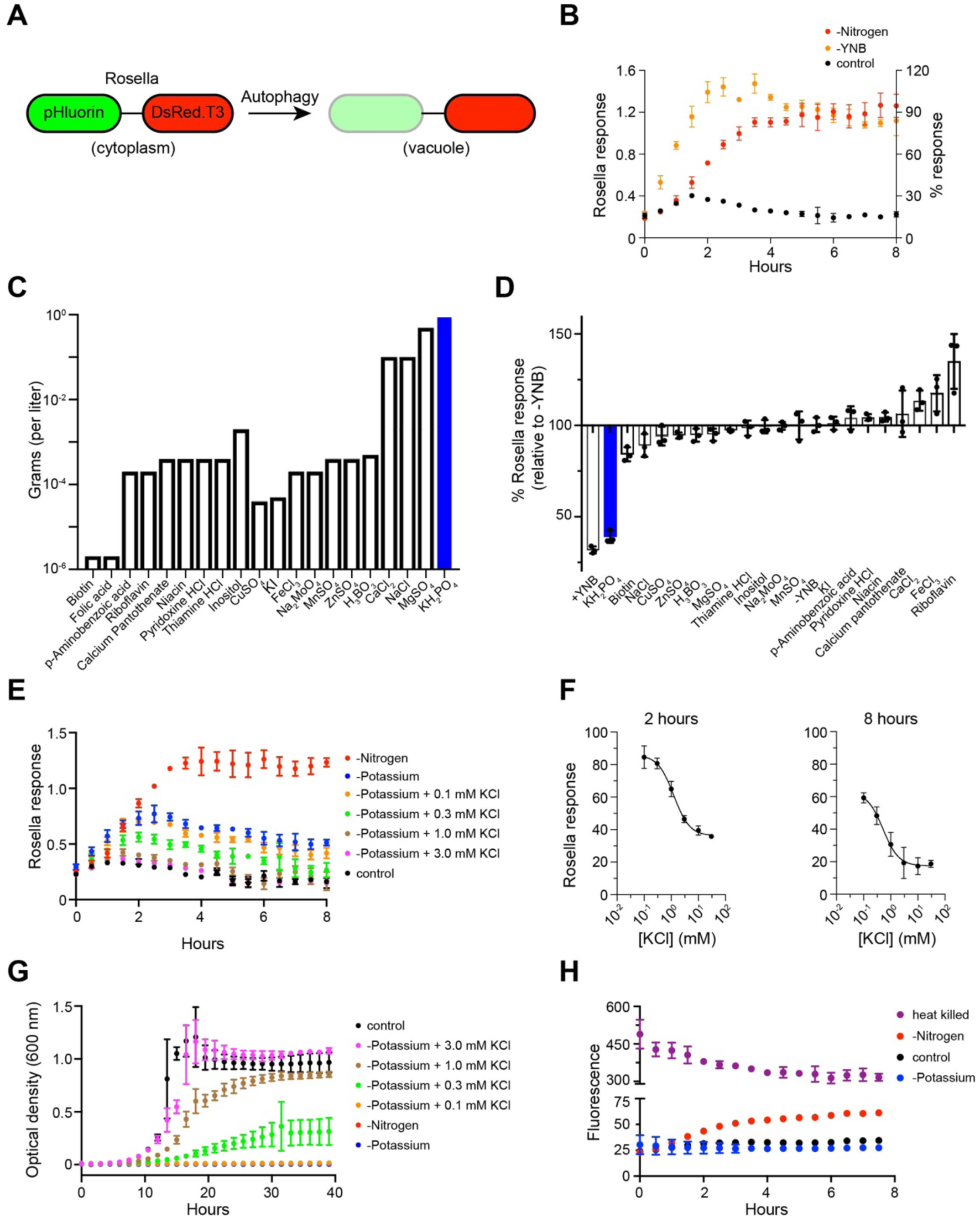
Potassium starvation promotes autophagy. (A) Rosella is comprised of super-ecliptic pHluorin (green) and DsRed.T3 (red). Upon induction of autophagy, Rosella is transported to the vacuole where low pH results in attenuation of green fluorescence, whereas red fluorescence is unaffected. (B) Cells expressing Rosella were maintained in exponential growth (OD at 600 nm <1) for 24 h prior to starvation with nitrogen or Yeast Nitrogen Base (YNB). Fluorescence was measured at 30 min intervals for 8 h. Response is presented as the ratio of red and green fluorescence. (C) Components of YNB and their abundance (in grams) in 1 liter of growth medium. (D) Cells were grown in SCD medium as described for (B), then transferred to SCD-YNB medium individually supplemented with each YNB component. Rosella response is shown as % relative to the response in SCD-YNB at 2 h. (E) Time-course of response in SCD-potassium medium upon addition of 0-3.0 mM KCl. (F) Dose-dependence of Rosella response obtained by plotting the 2 and 8 h data from (E). (G) Cellular growth under the same conditions as in (E), reported by optical density at 600 nm. (H) Time-course of cell viability measured as fluorescence from Propidium Iodide (PI). Rosella experiments were performed at least three times and data presented as ± standard deviation for four technical replicates. Growth and viability data are reported for three biological replicates. Data were analyzed in Microsoft Excel and GraphPad Prism.

Next, we used Rosella to test other nutrients important for proper cellular growth and homeostasis. As described in **Table 1**, <u>s</u>ynthetic <u>c</u>omplete medium with 2% <u>d</u>extrose (SCD) is composed of a nitrogen source (ammonium sulfate), sugar (dextrose), amino acids and nucleotides, as well as a base mixture (<u>y</u>east <u>n</u>itrogen <u>b</u>ase or YNB) comprised of vitamins, trace elements and salts. YNB does not provide nitrogen, which is included separately as ammonium sulfate. Other investigators have shown that sugar starvation leads to induction, but not completion, of autophagy (39,40). Therefore, we measured Rosella response in medium lacking YNB (SCD-YNB). As shown in **Figure 1B**, cells showed an increase in response that was similar in magnitude and substantially faster (t_1/2_ ∼1.25 h) than that observed in low nitrogen media (t_1/2_ ∼4.00 h). To identify the component(s) of YNB that individually contributed to the Rosella response (**Figure 1B**) we supplemented SCD-YNB medium with each individual vitamin, trace element or salt (**Table 2** and **Figure 1C**), and monitored fluorescence over time. As shown in **Figure 1D**, Rosella response was significantly diminished upon addition of potassium phosphate (+KH_2_PO_4_) alone, while readdition of other salts had no effect (**Figure S1**). We note the comparatively small response to zinc sulfate. Depletion for Zn^2+^ ions was reported to induce autophagy, but over longer timescales of >16 h (33). We conclude that, by this measure, potassium is the primary driver of autophagy observed in the absence of YNB.

**TABLE 1:** Components of Synthetic Complete medium with 2% Dextrose (SCD)

| Nutrient | Name | Amount (gram per liter, unless specified otherwise) |
| --- | --- | --- |
| Sugar | Dextrose | 20 |
| Nitrogen | Ammonium sulfate | 5 |
| Nucleotides + Amino acids | CSM Dropout mixture (CSM-Ura) | 0.69 |
|  | Uracil | 20 mg |
|  | Adenine | 30 mg |
|  | NaOH | 1 pellet |
| Vitamins, compounds supplying trace elements, and salts | Yeast Nitrogen Base (YNB) without amino acids and without ammonium sulfate | 1.7 |

**TABLE 2:** Components of Yeast Nitrogen Base (without amino acids and without ammonium sulfate)

| Nutrient | Name | Amount (microgram per liter) |
| --- | --- | --- |
| Vitamins | Biotin | 2 |
|  | Calcium pantothenate | 400 |
|  | Folic acid | 2 |
|  | Inositol | 2000 |
|  | Niacin | 400 |
|  | p-Aminobenzoic acid | 200 |
|  | Pyridoxine hydrochloride | 400 |
|  | Riboflavin | 200 |
|  | Thiamine hydrochloride | 400 |
| Compounds supplying trace elements | Boric acid | 500 |
|  | Copper sulfate | 40 |
|  | Potassium iodide | 100 |
|  | Ferric chloride | 200 |
|  | Manganese sulfate | 400 |
|  | Sodium molybdate | 200 |
|  | Zinc sulfate | 400 |
| Salts | Potassium phosphate | 1000 |
|  | Magnesium sulfate | 500 |
|  | Sodium chloride | 100 |
|  | Calcium chloride | 100 |

To distinguish between the contributions of the constituent ions (K^+^ and H_2_PO_4_^-^), we measured autophagy in cells exposed to growth medium in which potassium phosphate (in the YNB) was replaced with ammonium phosphate (SCD-potassium). As shown in **Figure 1E**, Rosella signal was elevated in these conditions, indicating that autophagy is specifically regulated by potassium cations. The response was recovered by the addition of 0.1-3 mM KCl, which has been used by others to replenish extracellular K^+^ ions (41,42). In complete SCD medium, K^+^ ions are present at a concentration of ∼7 mM. These concentrations are substantially lower than those required to activate the Hog1 kinase (>50 mM), which is crucial for cellular adaptation to external osmotic changes (43,44). As shown in the time profiles in **Figure 1E**, the response at 2 h was similar for both nitrogen and potassium, while the signals diverged substantially at later time points. The autophagy response at 8 h showed an EC_50_ of 1.24 mM KCl (**Figure 1F**).

Potassium ions are highly abundant within yeast cells (∼150-300 mM) and regulate ionic strength, turgor pressure, and enzyme function (28,45,46). In accordance with these requirements and as shown previously, cell growth is slowed by the lack of extracellular potassium (47). Addition of 0.3 mM KCl to SCD-potassium medium resulted in partial recovery of growth **(Figure 1G)**. At 1.0 and 3.0 mM KCl, long term recovery of growth was near complete. The growth response at 20 h showed an EC_50_ of 0.85 mM KCl **(Figure S2).** To ensure that cells remained viable, we monitored Propidium Iodide (PI) fluorescence (48) from individual cells in microplate wells exposed to the same growth media as in **Figure 1G**. As shown in **Figure 1H**, PI signal remained low under potassium starvation conditions and was elevated during nitrogen starvation. Thus, potassium is required for proper cell growth and autophagy. The autophagy response is approximately one third that seen in response to nitrogen limitation. In contrast to nitrogen however, potassium is not required to maintain cell viability, at least in the short term.

### Potassium-dependent autophagy is mediated by autophagosomes

Our findings presented in **Figure 1** indicate that the potassium-dependent autophagy response is four-fold lower than that for nitrogen. The magnitude of autophagy is regulated by many factors that control the number or size of autophagosomes (49,50). To further compare potassium and nitrogen starvation, we examined the accumulation of autophagosomes in individual cells using transmission electron microscopy (TEM). For these experiments, we used cells lacking the vacuolar protease Pep4, which is required for breakdown of autophagosomes within the vacuole (6). We first grew the cells for 6 h in SCD-nitrogen or SCD-potassium medium, harvested by centrifugation at 4000*g* for 1 min and prepared for TEM imaging (fixed, dehydrated, embedded, cut and stained). As expected, nitrogen starvation resulted in the formation of multiple autophagosomes in ∼95% of the cells. On average, we observed 6.9 ± 0.9 autophagosomes per cell with a cross-sectional area of 0.102 ± 0.042 μm^2^ (**Figures 2A and 2B**). In potassium-starved samples, autophagosomes were evident in only ∼10% of the cells. However, the number (6.5 ± 1.0 per cell) and size (0.089 ± 0.037 μm^2^) of autophagosomes was similar to the nitrogen response. In both conditions, we observed accumulation of glycogen granules (G) and lipid droplets (L) in the cytoplasm (**Figure 2A and 2C**). These features are characteristic of bulk autophagy (6). While most of the autophagosomes observed in our TEM images were already engulfed by the vacuole, we also observed some that were fused to the vacuolar membrane (**Figure 2C**) prior to engulfment. From these results, we conclude that potassium starvation leads to autophagosome formation, but does so in only a subset of cells.

**Figure 2:**
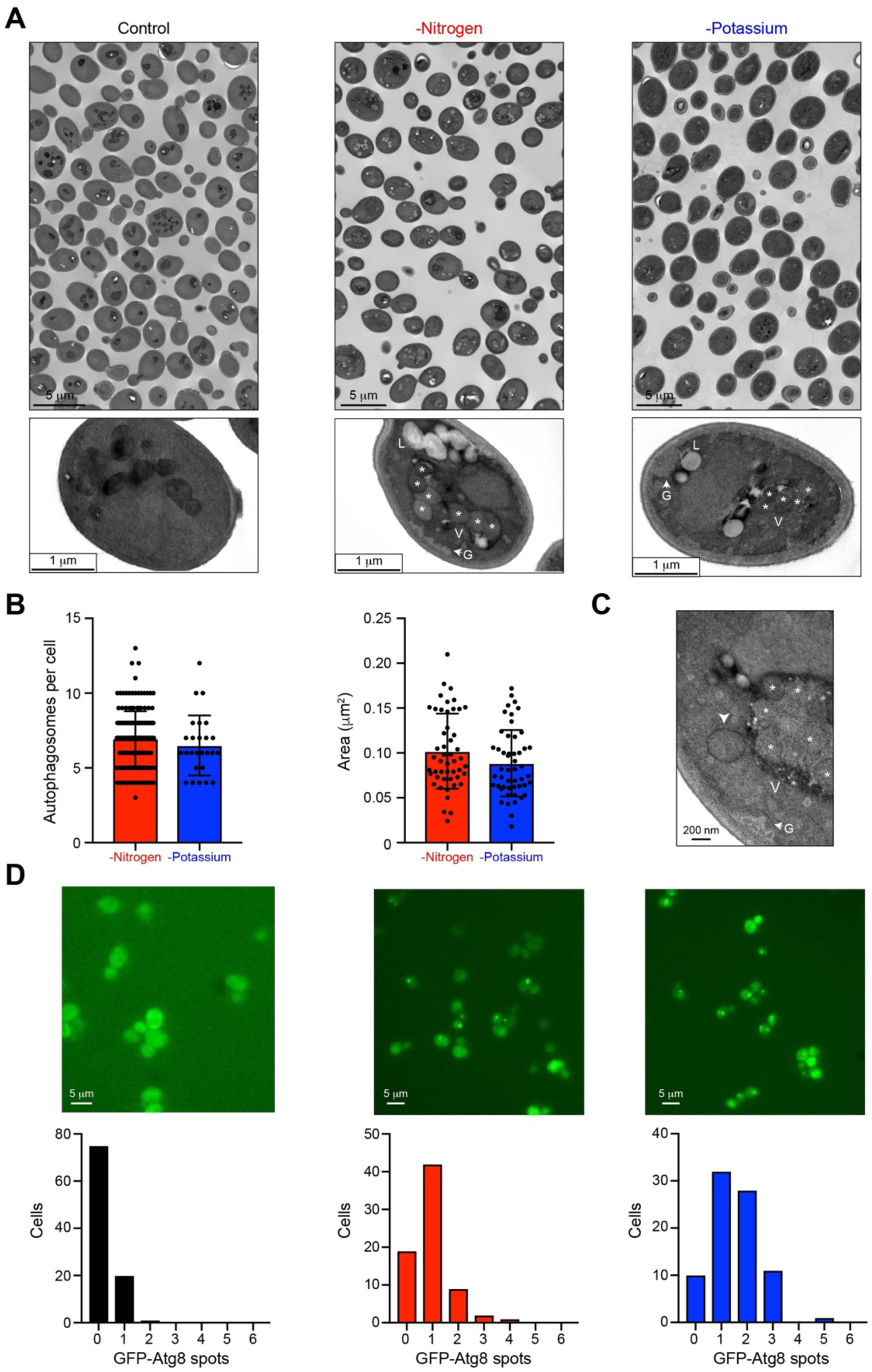
Potassium-dependent autophagy is mediated by autophagosomes. (A) Representative images of individual cell sections after 6 h of nitrogen or potassium starvation. Vacuole (V), autophagosomes (*), lipid droplet (L), glycogen (G). (B) Distribution of the abundance (per cell) and size (area in μm^2^) of autophagosomes (n = 50). (C) High magnification image (80,000x) showing an autophagosome (arrowhead) fused to the vacuolar membrane. (D) Representative single cell images and distributions of GFP-Atg8 spots formed in response to 2 h of nitrogen or potassium limitation (n = 85 cells).

Whereas Rosella reports on the transport of cytoplasmic cargo to the vacuole, other methods directly monitor components of the autophagy machinery such as autophagosomes and vacuolar proteins. To confirm the pro-autophagy effect of potassium starvation, we monitored the spatial localization of the ubiquitin-like protein Atg8. Upon induction of autophagy, Atg8 conjugates with phosphatidylethanolamine to form autophagosomes, which deliver cytoplasmic contents to the vacuole (19,51). During this process, Atg8 assembles into autophagosomal clusters, which are observed as bright puncta within the cell (52). As shown in **Figure 2D**, we observed a substantial increase in GFP-Atg8 clusters under potassium deprivation. This response was greater than that for nitrogen-starved cells, as is evident from the frequency distribution plots. These results indicate that autophagy is initiated in a majority of the cells, but the process is completed in only a subset of those cells.

### Potassium-dependent autophagy requires core ATG kinases and the PI 3-kinase Complex I

After being internalized by the vacuole, the autophagosomal cargo is released for degradation and recycling. This process can be monitored by cleavage of GFP-Atg8 (**Figure 3A**) (53). As shown in **Figure 3B**, GFP was observed after 3 h of potassium starvation and the response increased further at 6 h, indicating a sustained effect. We validated our findings from the Rosella and GFP-Atg8 methods using the Pho8Δ60 enzymatic assay (54). The alkaline phosphatase Pho8 is translocated to the vacuole using the secretory pathway. While the truncated form is maintained in the cytosol (**Figure 3C**), it is transported to the vacuole under low nitrogen conditions and subsequently activated via proteolytic processing. The increase in enzyme activity is a quantitative indicator of autophagy, and is measured as the conversion of a substrate to a fluorescent product. As shown in **Figure 3C** and in agreement with our findings from the other assays, potassium deprivation resulted in an increase in activity that was ∼one-third that observed with nitrogen starvation.

**Figure 3:**
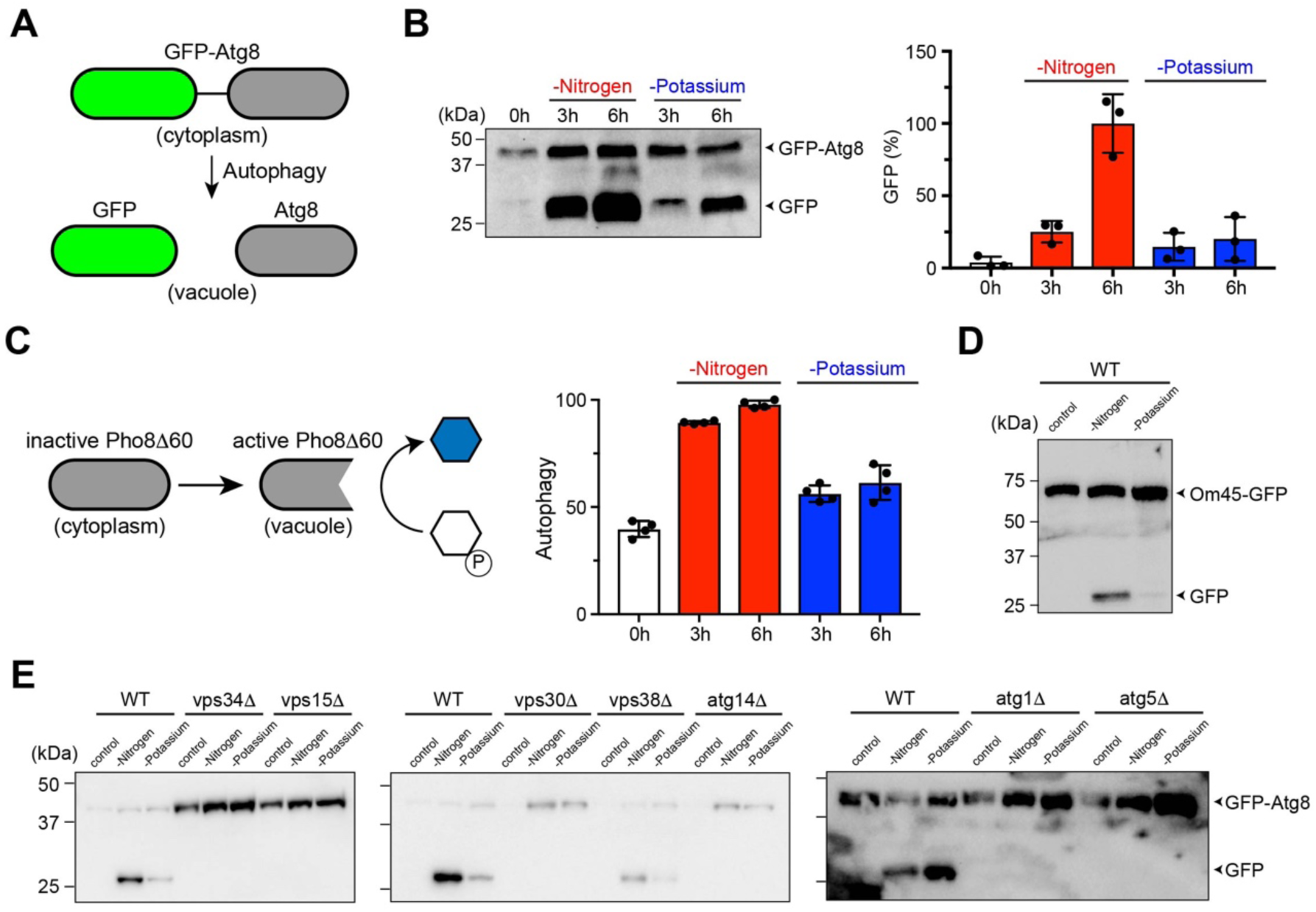
Potassium-dependent autophagy requires autophagy-related kinases Atg1 and Atg5, and the PI 3-kinase Complex I. (A) In pro-autophagy conditions, the GFP-Atg8 reporter is transported to the vacuole and enzymatically processed to release GFP. (B) GFP-Atg8 processing in wild-type BY4741 cells after 0, 3 and 6 h of starvation for nitrogen or potassium. Protein bands were quantified using ImageLab and presented as percentage of GFP-Atg8. (C) Induction of autophagy results in activation of the Pho8Δ60 enzyme, which converts a substrate into a fluorescent product. Shown are data for Pho8Δ60 activity in the same conditions as (B). (D) Processing of mitophagy reporter Om45-GFP in wild-type BY4741 after 6 h of nitrogen or potassium starvation. (E) Processing of autophagy reporter GFP-Atg8 after 6 h of starvation for nitrogen or potassium in BY4741 cells lacking Atg1, Atg5, Vps34, Vps15, Vps30, Vps38 or Atg14. All experiments were repeated three times and data are ± standard deviation for three or four biological replicates.

In addition to the bulk cytoplasm, organelles such as mitochondria, peroxisomes and ribosomes are also recognized as specific cargo for autophagic degradation. These selective forms of autophagy employ many of the core pathway components, as well as cargo-specific receptors (55-57). Previously, it was shown that pharmacological alterations to potassium ion homeostasis can lead to the production of reactive oxygen species, mitochondrial damage, and autophagic degradation of mitochondria (mitophagy) (58). Given these findings, we sought to determine the effects of our potassium limiting conditions on mitophagy. To that end, we monitored two GFP-tagged proteins, the mitochondrial outer membrane protein Om45 and the matrix protein Idh1 (59-61). As shown in **Figures 3D and S3**, whereas nitrogen starvation resulted in the release of GFP from these proteins, potassium starvation had a negligible effect. Thus, bulk autophagy is regulated by nitrogen or potassium whereas mitophagy is dependent on nitrogen alone.

Vps34 is the sole Phosphatidylinositol 3-kinase (PI3-kinase) in yeast and is essential for autophagy. Phosphatidylinositol 3-phosphate, generated by Vps34, is localized to autophagosomal membranes and is required for recruitment of other autophagy-related (ATG) proteins. Complex I is comprised of Vps34, a regulatory kinase Vps15, Vps30 (Beclin-1 in animal cells), and Atg14. Complex II contains Vps38 (UVRAG) in place of Atg14 (11-13). Whereas Complex I mediates autophagy in nutrient-limiting conditions, Complex II is essential for proper sorting of vacuolar enzymes under nutrient rich conditions (**Figure 3E**). To test the role of the effector complexes in potassium-dependent autophagy, we analyzed individual gene deletions using the GFP-Atg8 immunoblotting assay. As anticipated, we observed no processing of GFP-Atg8 in cells lacking Vps34 and Vps15 (**Figure 3E**). Similarly, autophagy was completely abrogated in cells lacking Atg14 (Complex I) or the common adapter Vps30, and only partially reduced by deletion of Vps38 (Complex II) (**Figures 3E and S4**). Potassium-dependent autophagy required the canonical components Atg1 and Atg5, in a manner similar to the nitrogen response (**Figure 3E**). This is consistent with recent work suggesting reciprocal regulation of potassium signaling and TORC1 activity, which inhibits autophagy by phosphorylating Atg13, a component of the Atg1 kinase complex (41,62). Thus, both potassium- and nitrogen-dependent autophagy are mediated by Atg1, Atg5 and the PI 3-kinase Complex I.

### Potassium and nitrogen starvation exhibit distinct transcriptional profiles

Our data indicate that potassium and nitrogen regulate a common autophagy response to supply raw materials for cellular biosynthesis. To determine how this information is interpreted, we first considered phosphorylation-mediated activation of the mitogen-activated protein kinase (MAPK) Kss1. In addition to its effects on autophagy, nitrogen depletion promotes cellular remodeling events such as filamentous growth (63), as well as the induction of genes involved in these processes, such as *FUS1* (64-67). Notably, filamentation requires elements of the MAPK pathway (68) and Kss1 activation is important for the transcription of filamentation genes under low nitrogen conditions (69). Preliminary evidence from others suggested that potassium depletion also induces *FUS1* transcription (42). However, the effect of nitrogen or potassium starvation on Kss1 activation has not been investigated and more generally, the regulatory mechanisms shared by the autophagy and filamentous growth pathways are not fully understood (70,71). To address this question, we used phos-tag gel electrophoresis to determine if either stimulus led to phosphorylation of Kss1. As expected, Kss1 was activated following depletion of nitrogen, although the effect was substantially less than that reported previously for addition of mating pheromone (**Figure S5**) (72). In contrast, Kss1 was unaffected by depletion of potassium.

Another way to understand the consequences of autophagy is to measure global changes in gene transcription. To that end, we sequenced the transcriptome of cells starved of potassium or nitrogen for 60 min (in duplicate) prior to RNA extraction and sequencing (**Figure 4A)**. Using principal component analysis (PCA), we observed that the two stimuli result in substantially different transcriptional profiles **(Figure S6)**. To further understand these differences, we carried out differential expression analysis, which revealed a dramatic impact on the transcriptome. Nitrogen starvation produced a highly dispersed response that included a large number of differentially expressed genes (DEGs) (log_2_(fold change) > 1 and adjusted p-value < 0.05) **(Figure 4B)**. In comparison, potassium starvation resulted in fewer DEGs with substantially smaller fold-change values. Using this approach, we identified 107 DEGs common to the two conditions **(Figure 4C)**. In addition to these shared targets, we found 68 and 626 unique DEGs for potassium and nitrogen, respectively. To understand the functional processes targeted by these stimuli, we performed gene ontology enrichment analysis using the DEGs independently for each condition. Whereas potassium targets processes that organize and modify cellular components and organelles, nitrogen affects transport of copper and transition metal ions **(Figure 5D)**. We conclude that potassium and nitrogen mediate their responses through distinct cellular mechanisms.

**Figure 4:**
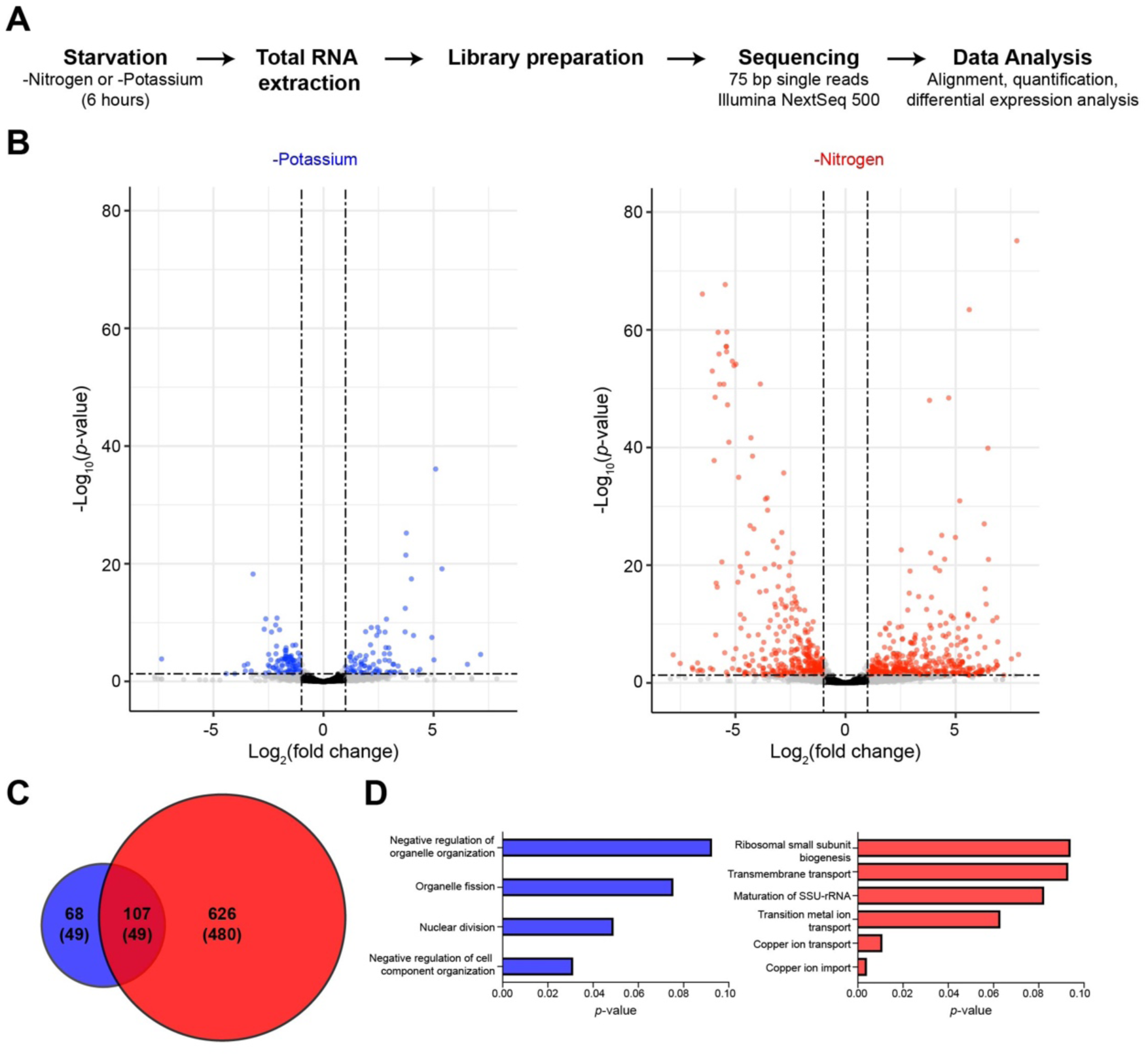
Potassium and nitrogen starvation exhibit distinct transcriptional profiles. (A) Schematic representation of experimental approach to obtain RNA sequencing data for potassium or nitrogen starvation. (B) Differential expression analysis reveals statistically significant changes in RNA abundance. Genes with adjusted p-value <0.05 and log_2_(fold change) > 1 are highlighted in color. (C) Venn diagram showing overlap between differentially expressed genes (DEGs) for potassium and nitrogen starvation. Numbers in parentheses represent genes with a known Gene Ontology annotation. (D) Gene Ontology process terms enriched amongst the DEGs for each condition. RNA abundance was quantified for two biological replicates for each condition.

## DISCUSSION

Most pioneering studies of autophagy use nitrogen starvation as the stimulus, primarily due to its large and rapid response. Here we demonstrate, using a panel of reporter assays in individual cells and populations, that potassium deprivation is a new and potent inducer of autophagy. The magnitude of the initial response is comparable to that observed with nitrogen starvation. Both stimuli transmit their effects via the canonical pathway components Atg1, Vps34, Vps15, Vps30, Atg14 and Atg5, which suggests a common route to cellular recycling. In contrast to these shared features, transcriptional responses to nitrogen and potassium starvation are substantially different. Collectively, our data indicate that nitrogen and potassium share the ability to supply metabolic building blocks, but adopt distinct ways to reallocate these resources for future use.

Traditional methods to study autophagy require extensive sample preparation prior to readout and are not conducive to high-throughput analysis. In comparison, the Rosella plate reader assay provides a convenient, automated platform for quantitative measurements in diverse genetic backgrounds and growth conditions. Our comprehensive analysis showed that while nitrogen transmits the largest autophagy effects, potassium is a close second. By integrating the Rosella reporter with other independent and well-established methods (electron microscopy, GFP-Atg8 immunoblotting and imaging, as well as the Pho8Δ60 alkaline phosphatase assay), we were able to compare nearly 30 conditions and multiple time points, thereby accelerating the discovery process.

Our findings add to a growing list of novel regulators of autophagy such as amino acids, sulfate, glucose and, most recently, zinc ions (6,36). The potassium response is robust and mirrors that of nitrogen limitation particularly at early time points (<2 h). This is in contrast with zinc, which induces a response that is comparatively small and is evident only after prolonged treatment (>16 h) (**Figure 1D** and (33)). Another class of inducers of autophagy operates through G protein coupled receptors (GPCRs). Prominent among these are the mammalian muscarinic cholinergic and β-adrenergic receptors, which bind to acetylcholine and epinephrine, respectively (73-78). Likewise, we have shown that the pheromone GPCR in yeast promotes vacuolar delivery of cytoplasmic contents (38,79). Whereas the response to nitrogen and potassium requires Complex I of the yeast PI 3-kinase, mating pheromone requires Complex II. Collectively, these studies demonstrate that multiple biochemically and physiologically distinct inputs converge on a common cellular process, one that is needed to support survival during changing growth conditions.

Although nitrogen and potassium induce a similar autophagy response, they are likely to serve distinct cellular needs. Nitrogen provides raw materials for synthesis of amino acids and nucleotides that enable the formation of proteins and nucleic acids. Potassium is accumulated within yeast and other eukaryotes to maintain important cellular parameters such as electroneutrality, pH, turgor pressure and cell volume. We observed that nitrogen targets genes that regulate transport of ions such as copper. In contrast, potassium impacts genes involved in a broader set of cellular processes, which overlap only partially with the nitrogen response. These data reveal that seemingly “redundant” input signals can nevertheless diverge and initiate distinct and complementary processes that prepare the cell for prolonged periods of nutrient deprivation.

In summary, our systematic and comprehensive studies highlight the power of high-throughput methods in hypothesis testing and discovery. Many important cellular pathways, including those mediated by the ATG machinery, were first discovered in yeast and subsequently confirmed in more complex organisms. Given the conservation of pathways governing autophagy and ion homeostasis, we expect that potassium likewise regulates autophagy-related processes in humans. Such mechanisms may be important in understanding the functional consequences of hypokalemia, as occurs in chronic kidney disease, alcohol abuse and diabetic ketoacidosis.

## EXPERIMENTAL PROCEDURES

### Strains, plasmids and growth media

Yeast strains used in this study were BY4741 (*MAT***a** *leu2*Δ *met15*Δ *his3*Δ *ura3*Δ), BY4741-derived gene deletion mutants obtained from the Yeast Knockout Collection (Invitrogen) or remade by homologous recombination of PCR-amplified drug resistance genes with flanking homology to the gene of interest (**Table S1**), and BY4741-derived GFP-gene fusions obtained from the Yeast GFP Clone Collection (Thermo Fisher Scientific) (80). *PHO8*Δ60 (gift from Mara Duncan, University of Michigan) was stably integrated into the genome, replacing the native *PHO8* gene. Plasmid pRS416-GFP-ATG8 (Addgene 49425) was a gift from Daniel Klionsky (81), and the GFP-*ATG8* construct was subsequently introduced into the pRS415 vector (82). Rosella plasmid pAS1NB-DsRed.T3-SEP (*2μ, amp*^*R*^, *LEU2*^*+*^) was a gift from Mark Prescott and Rodney Devenish (37). The Kss1– 9xMyc-tagged strain was generated by homologous recombination of a PCR-amplified 9xMyc cassette harboring a resistance gene to hygromycin B from plasmid pYM20 (pYM-9xMyc-hphNT1) at the C-terminus of the *KSS1* open reading frame (ORF) (83,84).

Cells were grown in rich medium containing yeast extract (10 gram/liter), peptone (20 gram/liter) and 2% (w/v) dextrose (YPD) or synthetic complete medium containing 2% (w/v) dextrose (SCD), ammonium sulfate, nucleotides, amino acids and a base mixture of vitamins, trace elements and salts (yeast nitrogen base, YNB) (**Table 1**). Plasmid selection was maintained by antibiotic supplementation or exclusion of appropriate nutrients. For nutrient starvation analysis, SCD-nitrogen medium was prepared by excluding ammonium sulfate, the major source of nitrogen. SCD-YNB medium was prepared by omitting yeast nitrogen base. To study individual YNB ingredients, SCD-YNB medium was supplemented with each component as listed in **Table 2**. Cells were cultured overnight with shaking at 30°C. Saturated cultures were diluted with fresh medium to optical density at 600 nm (OD_600_) = 0.1 and cultured for 6 h, then further diluted to OD_600_ = 0.001 and cultured for 18 h to maintain OD_600_ < 1 prior to use. For potassium starvation, cells were transferred to SCD-potassium medium, which was prepared using a modified YNB containing ammonium sulfate instead of potassium sulfate (Translucent K^+^ free medium containing ∼15 μM K^+^, ForMedium CYN7501). As a benchmark for autophagy, cells were transferred to low nitrogen medium (SCD-nitrogen) lacking ammonium sulfate.

### Rosella microplate-reader assay

Cells were transformed with pAS1NB-DsRed.T3-SEP and cultured in SCD-leucine medium. To start the starvation time-course, 1 ml of cells at OD_600_ ∼ 1 were transferred to SCD-potassium medium after centrifugation at 13,000*g* for 1 min, washing and resuspension. 200 μL of cells were then added to individual wells in black clear-bottom 96-well microplates (Greiner 655087 or Corning 3631). Nitrogen-starved cells served as a reference treatment for autophagy. Untreated ‘control’ cells in SCD-leucine medium were included in separate wells to enable measurement of basal response. Microplates were sealed to reduce evaporation (adhesive PCR plate seal, Thermo Fisher Scientific AB0558) and placed in a microplate reader (Molecular Devices SpectraMax i3x) for 8 h at 30°C. At each timepoint, samples were shaken and fluorescence was measured for super-ecliptic pHluorin (SEP) (488 nm excitation, 530 nm emission) and DsRed.T3 (543 nm excitation, 587 nm emission). Background signal was measured using clear, cell-free medium. All experiments were performed in duplicate with three technical replicates. For each time point, Rosella response was calculated as the ratio of background-corrected dsRed.T3 and SEP fluorescence. Starvation–induced response was normalized with basal response observed for cells maintained in SCD-leucine medium. Dose-response profiles were calculated for the 2 or 8 h time points using a variable slope (four parameters) nonlinear regression with least squares fit (GraphPad Prism).

### Growth and viability measurements

For cell growth measurements, wild-type BY4741 cells were cultured as described above to OD_600_ ∼1, then diluted to OD_600_ = 0.05 into either SCD, SCD-potassium or SCD-nitrogen medium. 200 μl cells were transferred to a 96-well microplate and OD_600_ was monitored in a SpectraMax i3x microplate reader at 30 min intervals for 48 h.

Cell viability was measured using a Celigo S Imaging Cytometer (Nexcelom) as described previously (83). Briefly, 200 μl of cells at OD_600_ ∼ 0.05 were mixed with 20 μM Propidium Iodide (P3566, Molecular Probes) and plated in half-area, black, clear-bottom 96-well plates (Greiner CELLSTAR) by centrifugation at 500*g* for 5 min at 4°C. Cells were imaged at 30 min intervals for 8 h at room temperature using the “Target 1+Mask” settings. Green fluorescence from Propidium Iodide (PI) was measured with the inbuilt GFP channel, while brightfield images were acquired simultaneously and used as masks for cell segmentation using Celigo’s native algorithm. Debris and cell clumps were excluded from the analysis by gating analysis based on the mean intensity and aspect ratio of PI. Background correction was performed by subtracting residual intensity from cell-free regions on the microplate. For data presentation, the mean PI intensity was averaged across 10,000 cells for each condition.

### GFP-Atg8 immunoblotting

For the GFP-Atg8 processivity assay, cells were propagated in SCD-leucine medium prior to nitrogen or potassium starvation (85). Cell extracts were obtained after 0, 3 and 6 h of starvation. Cells were lysed with TCA buffer (10 mM Tris-HCl pH 8.0, 10% (w/v) trichloroacetic acid (TCA), 25 mM NH_4_OAc, 1 mM Na_2_EDTA). Protein extracts were reconstituted in resuspension buffer (100 mM Tris-HCl, 3% (w/v) sodium dodecyl sulfate (SDS), pH 11.0), and protein concentration was determined using the Bio-Rad DC assay. Samples were normalized to 0.5-1 μg/μl with resuspension buffer and sample buffer (500 mM Tris-HCl, 20% (v/v) glycerol, 2% (w/v) SDS, 200 mM dithiothreitol, 0.01% (w/v) bromophenol blue, pH 8.5). Proteins were resolved on 10% SDS-PAGE gels, transferred to nitrocellulose membranes and detected by first incubating with blocking buffer (5% non-fat milk in TBST with 10 mM sodium azide) and, subsequently, immunoblotting with GFP antibodies (sc-999c, clone B-2, Santa Cruz Biotechnology) at 1:1,000 dilution in blocking buffer or glucose 6-phosphate dehydrogenase (G6PDH) antibodies (8866, Cell Signaling Technology, 1:1,000). Immunoreactive species were detected with antibodies conjugated with horseradish peroxidase (715-035-150, Jackson ImmunoResearch) at 1:10,000 using ECL-plus reagent (Life Technologies). Protein bands were quantified on a BioRad Chemidoc Touch Imaging System using Image Lab software version 6.0.1 (Bio-Rad).

### GFP-Atg8 microscopy

Clustering of GFP-Atg8 was measured using brightfield illumination on a Nikon Ti 2000 inverted microscope. Agar pads were prepared by dissolving 2% (w/v) agar in SCD-leucine, SCD-potassium or SCD-nitrogen medium and spotting (200 μL) onto a clear glass slide. Another slide was quickly placed on top and the agarose was allowed to solidify. After removing the top slide, 5 μL of starved cells were mounted on the pad and sealed with a clean glass coverslip as described previously (86). GFP-Atg8 within single cells was imaged by excitation at 488 nm and emission at 500-550 nm. GFP-Atg8 spots were counted in 85-100 individual cells for each condition using FIJI image analysis software and graphs were plotted with Graphpad Prism (87).

### Om45-GFP and Idh1-GFP immunoblotting

BY4741 cells expressing Om45-GFP or Idh1-GFP were sourced from the Yeast GFP Clone Collection (Thermo Fisher Scientific) and maintained OD_600_ < 1 in YPD medium for ∼24 h. To promote increased expression of the GFP-tagged proteins, cells were then diluted 100-fold with YPL medium (2% lactate instead of dextrose) and cultured for 12-16 h. Starvation for nitrogen or potassium was carried out for 6 h prior to protein extraction and immunoblotting analysis, as described above for GFP-Atg8.

### Pho8Δ60 enzymatic assay

The Pho8Δ60 assay was performed as described previously (54). Cells were cultured for 24 h in SCD medium to OD_600_ ∼ 1 prior to transfer to SCD-nitrogen or SCD-potassium medium. After 0, 3 or 6 h of starvation, 5 ml cells were collected by centrifugation at 3000*g* for 5 min. Cell pellets were resuspended in 200 μl cold assay buffer (250 mM Tris-HCl pH 9.0, 10 mM MgSO_4_ and 10 μM ZnSO_4_), mixed with ∼100 μl acid-washed glass beads (500 μm diameter) and lysed on an automatic vortex mixer (5 min at 4°C). The suspension was diluted with 50 μl assay buffer and cell debris were removed by centrifugation at 13,000*g* for 5 min. Protein concentration in the supernatant solution was determined with the BioRad DC protein assay. To measure enzyme activity, cell lysates were diluted 10-fold and incubated with substrate (5 mM α-naphthyl phosphate, Sigma-Aldrich N7255, dissolved in assay buffer) at 30°C for 20 min. The reaction was stopped by adding 1 mL of 2 M glycine-NaOH (pH 11.0) solution. Fluorescence was measured on a SpectraMax i3x microplate reader (345 nm excitation, 472 nm emission) and enzyme activity calculated as emission per unit protein (mg) in the reaction.

### Transmission electron microscopy

BY4741 cells lacking the Pep4 vacuolar protease were maintained at OD_600_ < 1 for ∼24 h prior to starvation for nitrogen or potassium for 6 h. Cells were collected by centrifugation (1000*g* for 2 min) and samples were prepared for transmission electron microscopy according to standard methods (88,89). Cells were fixed with 2% (w/v) glutaraldehyde in 0.1M PIPES buffer (pH 6.8 and containing 1 mM MgCl_2_, 1 mM CaCl_2_, and 0.1 M sorbitol) at room temperature for 1 h and stored at 4°C for several days. The cell wall material was permeabilized by mixing with 1% (w/v) sodium metaperiodate. Pellets were post-fixed with 1% osmium tetroxide/1.25% potassium ferrocyanide (w/v)/0.1 M PIPES buffer (pH 6.8) for 1 h and stained *en bloc* in 2% (v/v) aqueous uranyl acetate for 20 min. Cell pellets were dehydrated using a series of increasing ethanol concentrations, rinsed with 100% propylene oxide and embedded in Spurr’s epoxy resin (Electron Microscopy Sciences, Hatfield, PA). Ultrathin sections (70-80 nm) were cut with a diamond knife, mounted on 200 mesh copper grids and stained with 4% (v/v) aqueous uranyl acetate for 12 min and with Reynold’s lead citrate for 8 min (90). Samples were observed using a JEOL JEM-1230 transmission electron microscope operating at 80kV (JEOL USA, Inc., Peabody, MA) and images were acquired with a Gatan Orius SC1000 CCD Digital Camera and Gatan Microscopy Suite 3.0 software (Gatan, Inc., Pleasanton, CA).

TEM images were analyzed using FIJI image processing software (87). Autophagosome frequency (number per cell) was estimated by manual counting for 50 representative cells. To measure autophagosome size, outlines of individual autophagosomes were traced using the freehand selection tool. Next, the image was spatially calibrated using the ‘Set scale’ tool to convert pixels to micrometers. Area (in square micrometers) was measured using the ‘Measure’ tool within the region of interest (ROI) Manager. Lipid droplets and vesicles were excluded from the analysis based on their location (vacuole vs cytoplasm), brightness (lipid droplets were significantly brighter) and size (vesicles were 5-10 fold smaller) relative to autophagosomes.

### PhosphoMAPK analysis

MAP kinase activation was measured using quantitative immunoblotting as previously described (83). Briefly, BY4741 Kss1-9xMyc cells were grown to OD_600_ ∼ 1 in SCD medium and transferred to SCD-nitrogen or SCD-potassium medium (starvation conditions) or treated with 3 μM α factor pheromone (positive control). Aliquots were collected at the indicated time points, mixed with 5% (w/v) TCA, and collected by centrifugation at 4000*g* for 2 min. Cell pellets were resuspended in TCA buffer and protein extracts were normalized with resuspension buffer and sample buffer. Proteins were resolved at 150V for 1.5 h at 25°C with 10% SDS-PAGE gels containing 50 μM Phos-tag (Wako Chemicals) and 100 μM Zn(NO_3_)_2_. Proteins were then transferred to polyvinyledene difluoride (PVDF) membranes (Millipore IPVH00010) in Phos-tag transfer buffer at 20V for 20 h at 4°C. Blots were probed with Myc antibodies (2276, Cell Signaling Technology, 1:1000) or glucose 6-phosphate dehydrogenase (8866, Cell Signaling Technology, 1:1000) antibodies and quantified as detailed above.

### RNA sequencing

Cells were grown to OD_600_ ∼ 1 in SCD medium, then starved for nitrogen or potassium for 1 h. For each condition, 8×10^7^ cells were collected by centrifugation at 11,000*g* for 30 s at 4°C. The supernatant was aspirated and cell pellets were frozen in liquid nitrogen and stored at −80°C. Total RNA was extracted using the RNeasy Mini kit (Qiagen 74104) with on-column removal of DNA using the RNase-Free DNase Set (Qiagen 79254). The cDNA library was prepared with the KAPA mRNA HyperPrep Kit (Roche 08098115702, KAPA Code KK8580), barcoded with NEBNext Multiplex Oligos (Illumina E7710S), and sequenced with Illumina NextSeq 500 for 75bp single-end reads.

Quality check of raw sequence data was performed using the FastQC algorithm (91). Genome indices for yeast were downloaded from Ensembl.org and sequence alignment was performed using the STAR algorithm (92). Transcripts were quantified with the SALMON algorithm (93). Data were filtered to remove genes with <10 counts across all conditions and analyzed with DESeq2 package in R, while controlling for batch effect (94,95). All reported log_2_ fold-change values and adjusted p-values for genes were directly obtained from the DESeq2 model. GO term enrichment analysis was performed with the Gene Ontology Term Finder (version 0.86) from the Saccharomyces Genome Database (https://www.yeastgenome.org/).

## DATA AVAILABILITY STATEMENT

All the data described herein are presented within this manuscript.

## CONFLICT OF INTEREST

The authors declare that they have no conflicts of interest with the contents of this article. The content is solely the responsibility of the authors and does not necessarily represent the official views of the National Institutes of Health.

## FOOTNOTES

Research reported in the publication was supported by the National Institutes of Health under award number R35GM118105 (to H.G.D.). P.D. was supported by an individual Summer Undergraduate Research Fellowship (SURF) from the Office for Undergraduate Research at UNC Chapel Hill. The Microscopy Services Laboratory, Department of Pathology and Laboratory Medicine, is supported in part by P30 CA016086 National Cancer Institute (NCI) Center Core Support Grant to the UNC Lineberger Comprehensive Cancer Center. We thank Paul Cullen for valuable feedback on the manuscript, Lee Bardwell for helpful comments, Patrick Brennwald for help with interpretation of TEM images; and Sara Kimiko Suzuki and Shu Zhang for assistance with strain construction and immunoblotting.

## ABBREVIATIONS

The abbreviations used are:

PI: Phosphatidylinositol
ATG: Autophagy-related gene
SCD: Synthetic Complete medium with Dextrose
YNB: Yeast Nitrogen Base
TEM: Transmission Electron Microscopy
MAPK: Mitogen-Activated Protein Kinase
DEG: Differentially Expressed Gene.

## Notes

https://www.ncbi.nlm.nih.gov/geo/query/acc.cgi?acc=GSE151898

### Competing Interest Statement

The authors have declared no competing interest.

## REFERENCES

1. Jona, G., Choder, M., and Gileadi, O. (2000) Glucose starvation induces a drastic reduction in the rates of both transcription and degradation of mRNA in yeast. Biochim Biophys Acta 1491, 37–48

2. Lillie, S. H., and Pringle, J. R. (1980) Reserve carbohydrate metabolism in Saccharomyces cerevisiae: responses to nutrient limitation. J Bacteriol 143, 1384–1394

3. Martinez-Pastor, M. T., and Estruch, F. (1996) Sudden depletion of carbon source blocks translation, but not transcription, in the yeast Saccharomyces cerevisiae. FEBS Lett 390, 319–322

4. Nilsson, A., Pahlman, I. L., Jovall, P. A., Blomberg, A., Larsson, C., and Gustafsson, L. (2001) The catabolic capacity of Saccharomyces cerevisiae is preserved to a higher extent during carbon compared to nitrogen starvation. Yeast 18, 1371–1381

5. Dikic, I. (2017) Proteasomal and Autophagic Degradation Systems. Annu Rev Biochem 86, 193–224

6. Takeshige, K., Baba, M., Tsuboi, S., Noda, T., and Ohsumi, Y. (1992) Autophagy in yeast demonstrated with proteinase-deficient mutants and conditions for its induction. J Cell Biol 119, 301–311

7. Mitchener, J. S., Shelburne, J. D., Bradford, W. D., and Hawkins, H. K. (1976) Cellular autophagocytosis induced by deprivation of serum and amino acids in HeLa cells. Am J Pathol 83, 485–492

8. Schmelzle, T., and Hall, M. N. (2000) TOR, a central controller of cell growth. Cell 103, 253–262

9. De Virgilio, C., and Loewith, R. (2006) Cell growth control: little eukaryotes make big contributions. Oncogene 25, 6392–6415

10. Cebollero, E., and Reggiori, F. (2009) Regulation of autophagy in yeast Saccharomyces cerevisiae. Biochim Biophys Acta 1793, 1413–1421

11. Schu, P. V., Takegawa, K., Fry, M. J., Stack, J. H., Waterfield, M. D., and Emr, S. D. (1993) Phosphatidylinositol 3-kinase encoded by yeast VPS34 gene essential for protein sorting. Science 260, 88–91

12. Stack, J. H., Herman, P. K., Schu, P. V., and Emr, S. D. (1993) A membrane-associated complex containing the Vps15 protein kinase and the Vps34 PI 3-kinase is essential for protein sorting to the yeast lysosome-like vacuole. Embo J 12, 2195–2204

13. Backer, J. M. (2016) The intricate regulation and complex functions of the Class III phosphoinositide 3-kinase Vps34. Biochem J 473, 2251–2271

14. Herman, P. K., Stack, J. H., and Emr, S. D. (1991) A genetic and structural analysis of the yeast Vps15 protein kinase: evidence for a direct role of Vps15p in vacuolar protein delivery. Embo J 10, 4049–4060

15. Stack, J. H., DeWald, D. B., Takegawa, K., and Emr, S. D. (1995) Vesicle-mediated protein transport: regulatory interactions between the Vps15 protein kinase and the Vps34 PtdIns 3-kinase essential for protein sorting to the vacuole in yeast. J Cell Biol 129, 321–334

16. Kametaka, S., Okano, T., Ohsumi, M., and Ohsumi, Y. (1998) Apg14p and Apg6/Vps30p form a protein complex essential for autophagy in the yeast, Saccharomyces cerevisiae. J Biol Chem 273, 22284–22291

17. Kihara, A., Noda, T., Ishihara, N., and Ohsumi, Y. (2001) Two distinct Vps34 phosphatidylinositol 3-kinase complexes function in autophagy and carboxypeptidase Y sorting in Saccharomyces cerevisiae. J Cell Biol 152, 519–530.

18. Yuan, H. X., Russell, R. C., and Guan, K. L. (2013) Regulation of PIK3C3/VPS34 complexes by MTOR in nutrient stress-induced autophagy. Autophagy 9, 1983–1995

19. Ichimura, Y., Kirisako, T., Takao, T., Satomi, Y., Shimonishi, Y., Ishihara, N., Mizushima, N., Tanida, I., Kominami, E., Ohsumi, M., Noda, T., and Ohsumi, Y. (2000) A ubiquitin-like system mediates protein lipidation. Nature 408, 488–492

20. Abeliovich, H., Dunn, W. A., Jr., Kim, J., and Klionsky, D. J. (2000) Dissection of autophagosome biogenesis into distinct nucleation and expansion steps. J Cell Biol 151, 1025–1034

21. Fujita, N., Itoh, T., Omori, H., Fukuda, M., Noda, T., and Yoshimori, T. (2008) The Atg16L complex specifies the site of LC3 lipidation for membrane biogenesis in autophagy. Mol Biol Cell 19, 2092–2100

22. Xie, Z., Nair, U., and Klionsky, D. J. (2008) Atg8 controls phagophore expansion during autophagosome formation. Mol Biol Cell 19, 3290–3298

23. Kaufmann, A., Beier, V., Franquelim, H. G., and Wollert, T. (2014) Molecular mechanism of autophagic membrane-scaffold assembly and disassembly. Cell 156, 469–481

24. Weidberg, H., Shvets, E., Shpilka, T., Shimron, F., Shinder, V., and Elazar, Z. (2010) LC3 and GATE-16/GABARAP subfamilies are both essential yet act differently in autophagosome biogenesis. EMBO J 29, 1792–1802

25. Lee, Y. K., and Lee, J. A. (2016) Role of the mammalian ATG8/LC3 family in autophagy: differential and compensatory roles in the spatiotemporal regulation of autophagy. BMB Rep 49, 424–430

26. Rubinsztein, D. C., Shpilka, T., and Elazar, Z. (2012) Mechanisms of autophagosome biogenesis. Curr Biol 22, R29–34

27. Merchan, S., Bernal, D., Serrano, R., and Yenush, L. (2004) Response of the Saccharomyces cerevisiae Mpk1 mitogen-activated protein kinase pathway to increases in internal turgor pressure caused by loss of Ppz protein phosphatases. Eukaryot Cell 3, 100–107

28. Page, M. J., and Di Cera, E. (2006) Role of Na+ and K+ in enzyme function. Physiol Rev 86, 1049–1092

29. Lubin, M., and Ennis, H. L. (1964) On the Role of Intracellular Potassium in Protein Synthesis. Biochim Biophys Acta 80, 614–631

30. Mancias, J. D., Pontano Vaites, L., Nissim, S., Biancur, D. E., Kim, A. J., Wang, X., Liu, Y., Goessling, W., Kimmelman, A. C., and Harper, J. W. (2015) Ferritinophagy via NCOA4 is required for erythropoiesis and is regulated by iron dependent HERC2-mediated proteolysis. Elife 4

31. Medina, D. L., and Ballabio, A. (2015) Lysosomal calcium regulates autophagy. Autophagy 11, 970–971

32. Medina, D. L., Di Paola, S., Peluso, I., Armani, A., De Stefani, D., Venditti, R., Montefusco, S., Scotto-Rosato, A., Prezioso, C., Forrester, A., Settembre, C., Wang, W., Gao, Q., Xu, H., Sandri, M., Rizzuto, R., De Matteis, M. A., and Ballabio, A. (2015) Lysosomal calcium signalling regulates autophagy through calcineurin and TFEB. Nat Cell Biol 17, 288–299

33. Kawamata, T., Horie, T., Matsunami, M., Sasaki, M., and Ohsumi, Y. (2017) Zinc starvation induces autophagy in yeast. J Biol Chem 292, 8520–8530

34. Pottier, M., Masclaux-Daubresse, C., Yoshimoto, K., and Thomine, S. (2014) Autophagy as a possible mechanism for micronutrient remobilization from leaves to seeds. Front Plant Sci 5, 11

35. Horie, T., Kawamata, T., Matsunami, M., and Ohsumi, Y. (2017) Recycling of iron via autophagy is critical for the transition from glycolytic to respiratory growth. J Biol Chem 292, 8533–8543

36. Sutter, B. M., Wu, X., Laxman, S., and Tu, B. P. (2013) Methionine inhibits autophagy and promotes growth by inducing the SAM-responsive methylation of PP2A. Cell 154, 403–415

37. Rosado, C. J., Mijaljica, D., Hatzinisiriou, I., Prescott, M., and Devenish, R. J. (2008) Rosella: a fluorescent pH-biosensor for reporting vacuolar turnover of cytosol and organelles in yeast. Autophagy 4, 205–213

38. Rangarajan, N., Gordy, C. L., Askew, L., Bevill, S. M., Elston, T. C., Errede, B., Hurst, J. H., Kelley, J. B., Sheetz, J. B., Suzuki, S. K., Valentin, N. H., Young, E., and Dohlman, H. G. (2019) Systematic analysis of F-box proteins reveals a new branch of the yeast mating pathway. J Biol Chem 294, 14717–14731

39. Lang, M. J., Martinez-Marquez, J. Y., Prosser, D. C., Ganser, L. R., Buelto, D., Wendland, B., and Duncan, M. C. (2014) Glucose starvation inhibits autophagy via vacuolar hydrolysis and induces plasma membrane internalization by down-regulating recycling. J Biol Chem 289, 16736–16747

40. Adachi, A., Koizumi, M., and Ohsumi, Y. (2017) Autophagy induction under carbon starvation conditions is negatively regulated by carbon catabolite repression. J Biol Chem 292, 19905–19918

41. Primo, C., Ferri-Blazquez, A., Loewith, R., and Yenush, L. (2017) Reciprocal Regulation of Target of Rapamycin Complex 1 and Potassium Accumulation. J Biol Chem 292, 563–574

42. Barreto, L., Canadell, D., Valverde-Saubi, D., Casamayor, A., and Arino, J. (2012) The short-term response of yeast to potassium starvation. Environ Microbiol 14, 3026–3042

43. Brewster, J. L., de Valoir, T., Dwyer, N. D., Winter, E., and Gustin, M. C. (1993) An osmosensing signal transduction pathway in yeast. Science 259, 1760–1763.

44. English, J. G., Shellhammer, J. P., Malahe, M., McCarter, P. C., Elston, T. C., and Dohlman, H. G. (2015) MAPK feedback encodes a switch and timer for tunable stress adaptation in yeast. Sci Signal 8, ra5

45. Yenush, L., Merchan, S., Holmes, J., and Serrano, R. (2005) pH-Responsive, posttranslational regulation of the Trk1 potassium transporter by the type 1-related Ppz1 phosphatase. Mol Cell Biol 25, 8683–8692

46. Haro, R., and Rodriguez-Navarro, A. (2002) Molecular analysis of the mechanism of potassium uptake through the TRK1 transporter of Saccharomyces cerevisiae. Biochim Biophys Acta 1564, 114–122

47. Lauff, D. B., and Santa-Maria, G. E. (2010) Potassium deprivation is sufficient to induce a cell death program in Saccharomyces cerevisiae. FEMS Yeast Res 10, 497–507

48. Deere, D., Shen, J., Vesey, G., Bell, P., Bissinger, P., and Veal, D. (1998) Flow cytometry and cell sorting for yeast viability assessment and cell selection. Yeast 14, 147–160

49. du Toit, A., Hofmeyr, J. S., Gniadek, T. J., and Loos, B. (2018) Measuring autophagosome flux. Autophagy 14, 1060–1071

50. Backues, S. K., Chen, D., Ruan, J., Xie, Z., and Klionsky, D. J. (2014) Estimating the size and number of autophagic bodies by electron microscopy. Autophagy 10, 155–164

51. Sugawara, K., Suzuki, N. N., Fujioka, Y., Mizushima, N., Ohsumi, Y., and Inagaki, F. (2004) The crystal structure of microtubule-associated protein light chain 3, a mammalian homologue of Saccharomyces cerevisiae Atg8. Genes Cells 9, 611–618

52. Klionsky, D. J. and co-workers (2016) Guidelines for the use and interpretation of assays for monitoring autophagy (3rd edition). Autophagy 12, 1–222

53. Cheong, H., and Klionsky, D. J. (2008) Biochemical methods to monitor autophagy-related processes in yeast. Methods Enzymol 451, 1–26

54. Noda, T., and Klionsky, D. J. (2008) The quantitative Pho8Delta60 assay of nonspecific autophagy. Methods Enzymol 451, 33–42

55. Kanki, T., Kang, D., and Klionsky, D. J. (2009) Monitoring mitophagy in yeast: the Om45-GFP processing assay. Autophagy 5, 1186–1189

56. Eberhart, T., and Kovacs, W. J. (2018) Pexophagy in yeast and mammals: an update on mysteries. Histochem Cell Biol 150, 473–488

57. Wang, X., Wang, P., Zhang, Z., Farre, J. C., Li, X., Wang, R., Xia, Z., Subramani, S., and Ma, C. (2020) The autophagic degradation of cytosolic pools of peroxisomal proteins by a new selective pathway. Autophagy 16, 154–166

58. Klein, B., Worndl, K., Lutz-Meindl, U., and Kerschbaum, H. H. (2011) Perturbation of intracellular K(+) homeostasis with valinomycin promotes cell death by mitochondrial swelling and autophagic processes. Apoptosis 16, 1101–1117

59. Kanki, T., and Klionsky, D. J. (2008) Mitophagy in yeast occurs through a selective mechanism. J Biol Chem 283, 32386–32393

60. Kanki, T., and Klionsky, D. J. (2009) Atg32 is a tag for mitochondria degradation in yeast. Autophagy 5, 1201–1202

61. Okamoto, K., Kondo-Okamoto, N., and Ohsumi, Y. (2009) Mitochondria-anchored receptor Atg32 mediates degradation of mitochondria via selective autophagy. Dev Cell 17, 87–97

62. Kamada, Y., Yoshino, K., Kondo, C., Kawamata, T., Oshiro, N., Yonezawa, K., and Ohsumi, Y. (2010) Tor directly controls the Atg1 kinase complex to regulate autophagy. Mol Cell Biol 30, 1049–1058

63. Gimeno, C. J., Ljungdahl, P. O., Styles, C. A., and Fink, G. R. (1992) Unipolar cell divisions in the yeast S. cerevisiae lead to filamentous growth: regulation by starvation and RAS. Cell 68, 1077–1090

64. Breitkreutz, A., Boucher, L., and Tyers, M. (2001) MAPK specificity in the yeast pheromone response independent of transcriptional activation. Curr Biol 11, 1266–1271.

65. Roberts, C. J., Nelson, B., Marton, M. J., Stoughton, R., Meyer, M. R., Bennett, H. A., He, Y. D., Dai, H., Walker, W. L., Hughes, T. R., Tyers, M., Boone, C., and Friend, S. H. (2000) Signaling and circuitry of multiple MAPK pathways revealed by a matrix of global gene expression profiles. Science 287, 873–880

66. Zeitlinger, J., Simon, I., Harbison, C. T., Hannett, N. M., Volkert, T. L., Fink, G. R., and Young, R. A. (2003) Program-specific distribution of a transcription factor dependent on partner transcription factor and MAPK signaling. Cell 113, 395–404

67. Paliwal, S., Iglesias, P. A., Campbell, K., Hilioti, Z., Groisman, A., and Levchenko, A. (2007) MAPK-mediated bimodal gene expression and adaptive gradient sensing in yeast. Nature 446, 46–51

68. Liu, H., Styles, C. A., and Fink, G. R. (1993) Elements of the yeast pheromone response pathway required for filamentous growth of diploids. Science 262, 1741–1744

69. Kusari, A. B., Molina, D. M., Sabbagh, W., Jr., Lau, C. S., and Bardwell, L. (2004) A conserved protein interaction network involving the yeast MAP kinases Fus3 and Kss1. J Cell Biol 164, 267–277

70. Ma, J., Jin, R., Jia, X., Dobry, C. J., Wang, L., Reggiori, F., Zhu, J., and Kumar, A. (2007) An interrelationship between autophagy and filamentous growth in budding yeast. Genetics 177, 205–214

71. Bruckner, S., Kern, S., Birke, R., Saugar, I., Ulrich, H. D., and Mosch, H. U. (2011) The TEA transcription factor Tec1 links TOR and MAPK pathways to coordinate yeast development. Genetics 189, 479–494

72. Nagiec, M. J., McCarter, P. C., Kelley, J. B., Dixit, G., Elston, T. C., and Dohlman, H. G. (2015) Signal inhibition by a dynamically regulated pool of monophosphorylated MAPK. Mol Biol Cell 26, 3359–3371

73. You, Y. J., Kim, J., Cobb, M., and Avery, L. (2006) Starvation activates MAP kinase through the muscarinic acetylcholine pathway in Caenorhabditis elegans pharynx. Cell Metab 3, 237–245

74. Oh, D. Y., Talukdar, S., Bae, E. J., Imamura, T., Morinaga, H., Fan, W., Li, P., Lu, W. J., Watkins, S. M., and Olefsky, J. M. (2010) GPR120 is an omega-3 fatty acid receptor mediating potent anti-inflammatory and insulin-sensitizing effects. Cell 142, 687–698

75. Aranguiz-Urroz, P., Canales, J., Copaja, M., Troncoso, R., Vicencio, J. M., Carrillo, C., Lara, H., Lavandero, S., and Diaz-Araya, G. (2011) Beta(2)-adrenergic receptor regulates cardiac fibroblast autophagy and collagen degradation. Biochim Biophys Acta 1812, 23–31

76. Lizaso, A., Tan, K. T., and Lee, Y. H. (2013) beta-adrenergic receptor-stimulated lipolysis requires the RAB7-mediated autolysosomal lipid degradation. Autophagy 9, 1228–1243

77. Wang, L., Lu, K., Hao, H., Li, X., Wang, J., Wang, K., Wang, J., Yan, Z., Zhang, S., Du, Y., and Liu, H. (2013) Decreased autophagy in rat heart induced by anti-beta1-adrenergic receptor autoantibodies contributes to the decline in mitochondrial membrane potential. PLoS One 8, e81296

78. Wauson, E. M., Zaganjor, E., Lee, A. Y., Guerra, M. L., Ghosh, A. B., Bookout, A. L., Chambers, C. P., Jivan, A., McGlynn, K., Hutchison, M. R., Deberardinis, R. J., and Cobb, M. H. (2012) The G protein-coupled taste receptor T1R1/T1R3 regulates mTORC1 and autophagy. Mol Cell 47, 851–862

79. Slessareva, J. E., Routt, S. M., Temple, B., Bankaitis, V. A., and Dohlman, H. G. (2006) Activation of the phosphatidylinositol 3-kinase Vps34 by a G protein alpha subunit at the endosome. Cell 126, 191–203

80. Huh, W. K., Falvo, J. V., Gerke, L. C., Carroll, A. S., Howson, R. W., Weissman, J. S., and O’Shea, E. K. (2003) Global analysis of protein localization in budding yeast. Nature 425, 686–691

81. Guan, J., Stromhaug, P. E., George, M. D., Habibzadegah-Tari, P., Bevan, A., Dunn, W. A., Jr., and Klionsky, D. J. (2001) Cvt18/Gsa12 is required for cytoplasm-to-vacuole transport, pexophagy, and autophagy in Saccharomyces cerevisiae and Pichia pastoris. Mol Biol Cell 12, 3821–3838

82. Sikorski, R. S., and Hieter, P. (1989) A system of shuttle vectors and yeast host strains designed for efficient manipulation of DNA in Saccharomyces cerevisiae. Genetics 122, 19–27

83. Shellhammer, J. P., Pomeroy, A. E., Li, Y., Dujmusic, L., Elston, T. C., Hao, N., and Dohlman, H. G. (2019) Quantitative analysis of the yeast pheromone pathway. Yeast 36, 495–518

84. Janke, C., Magiera, M. M., Rathfelder, N., Taxis, C., Reber, S., Maekawa, H., Moreno-Borchart, A., Doenges, G., Schwob, E., Schiebel, E., and Knop, M. (2004) A versatile toolbox for PCR-based tagging of yeast genes: new fluorescent proteins, more markers and promoter substitution cassettes. Yeast 21, 947–962

85. Shintani, T., and Klionsky, D. J. (2004) Cargo proteins facilitate the formation of transport vesicles in the cytoplasm to vacuole targeting pathway. J Biol Chem 279, 29889–29894

86. Tanaka, K., Kitamura, E., and Tanaka, T. U. (2010) Live-cell analysis of kinetochore-microtubule interaction in budding yeast. Methods 51, 206–213

87. Schindelin, J., Arganda-Carreras, I., Frise, E., Kaynig, V., Longair, M., Pietzsch, T., Preibisch, S., Rueden, C., Saalfeld, S., Schmid, B., Tinevez, J. Y., White, D. J., Hartenstein, V., Eliceiri, K., Tomancak, P., and Cardona, A. (2012) Fiji: an open-source platform for biological-image analysis. Nat Methods 9, 676–682

88. Vermulst, M., Denney, A. S., Lang, M. J., Hung, C. W., Moore, S., Moseley, M. A., Thompson, J. W., Madden, V., Gauer, J., Wolfe, K. J., Summers, D. W., Schleit, J., Sutphin, G. L., Haroon, S., Holczbauer, A., Caine, J., Jorgenson, J., Cyr, D., Kaeberlein, M., Strathern, J. N., Duncan, M. C., and Erie, D. A. (2015) Transcription errors induce proteotoxic stress and shorten cellular lifespan. Nat Commun 6, 8065

89. Wright, R. (2000) Transmission electron microscopy of yeast. Microsc Res Tech 51, 496–510

90. Reynolds, E. S. (1963) The use of lead citrate at high pH as an electron-opaque stain in electron microscopy. J Cell Biol 17, 208–212

91. Andrews, S. (2013) FastQC: A quality control tool for high throughput sequence data. Available online at: http://www.bioinformatics.babraham.ac.uk/projects/fastqc/

92. Dobin, A., Davis, C. A., Schlesinger, F., Drenkow, J., Zaleski, C., Jha, S., Batut, P., Chaisson, M., and Gingeras, T. R. (2013) STAR: ultrafast universal RNA-seq aligner. Bioinformatics 29, 15–21

93. Patro, R., Duggal, G., Love, M. I., Irizarry, R. A., and Kingsford, C. (2017) Salmon provides fast and bias-aware quantification of transcript expression. Nat Methods 14, 417–419

94. R Core Team (2013) R: A language and environment for statistical computing. R Foundation for Statistical Computing, Austria, Vienna. Available online at: http://www.R-project.org

95. Love, M. I., Huber, W., and Anders, S. (2014) Moderated estimation of fold change and dispersion for RNA-seq data with DESeq2. Genome Biol 15, 550

