## Supplemental Information for "Potassium starvation induces autophagy in yeast"

### Undergraduate researcher

*** To whom correspondence should be addressed:** Henrik G. Dohlman, University of North Carolina at Chapel Hill, 4016 Genetic Medicine Building, 120 Mason Farm Rd., Chapel Hill, NC 27599 USA. Telephone: 919-843-6894;

#### TABLE OF CONTENTS

**Figure S1**: Addition of $KH_2PO_4$ uniquely rescues autophagy in YNB-free medium.

**Figure S2:** Addition of KCl rescues growth inhibition in SCD-potassium medium.

**Figure S3:** Mitophagy is regulated by nitrogen, but not potassium.

**Figure S4:** Potassium dependent autophagy requires the PI 3-kinase Complex I.

**Figure S5:** Nitrogen starvation activates the MAP kinase Kss1.

**Figure S6:** Potassium and nitrogen starvation exhibit distinct transcriptional profiles.

**Table S1**: Yeast strains used in this study.

**FIGURE S1**

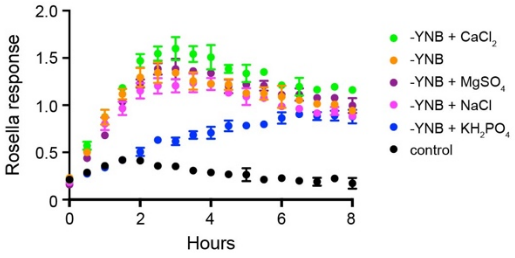


**Figure S1**: **$KH_2PO_4$ uniquely rescues autophagy in YNB-free medium.** Time-course of Rosella response in growth medium lacking YNB (SCD-YNB) after the individual readdition of each major salt component.

**FIGURE S2**

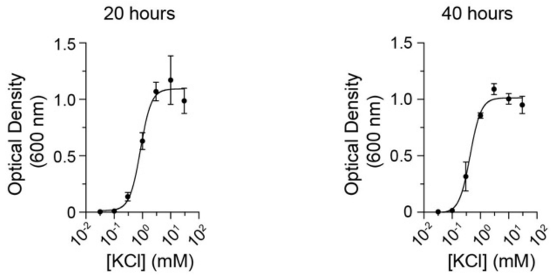


**Figure S2**: **Addition of KCl rescues growth inhibition in SCD-potassium medium.** Dose-dependence of cell growth reported by optical density at 600 nm ($OD_{600}$) at 20 h and 40 h (from **Figure 1G**).

**FIGURE S3**

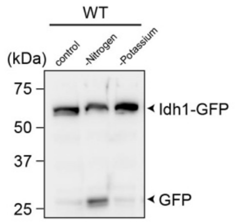


**Figure S3**: **Mitophagy is regulated by nitrogen, but not potassium.** Processing of mitophagy reporter Idh1-GFP in wild-type BY4741 after 6 h of nitrogen or potassium starvation. Data are representative of three biological replicates.

**FIGURE S4**

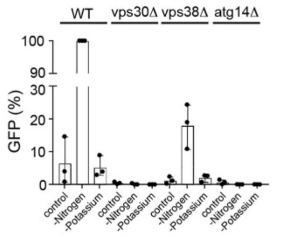


**Figure S4: Potassium dependent autophagy requires the PI 3-kinase Complex I.** Quantification of immunoblotting data on GFP-Atg8 processivity in BY4741 cells lacking Vps30 (Complex I and II), Vps38 (Complex II) or Atg14 (Complex I).

**FIGURE S5**

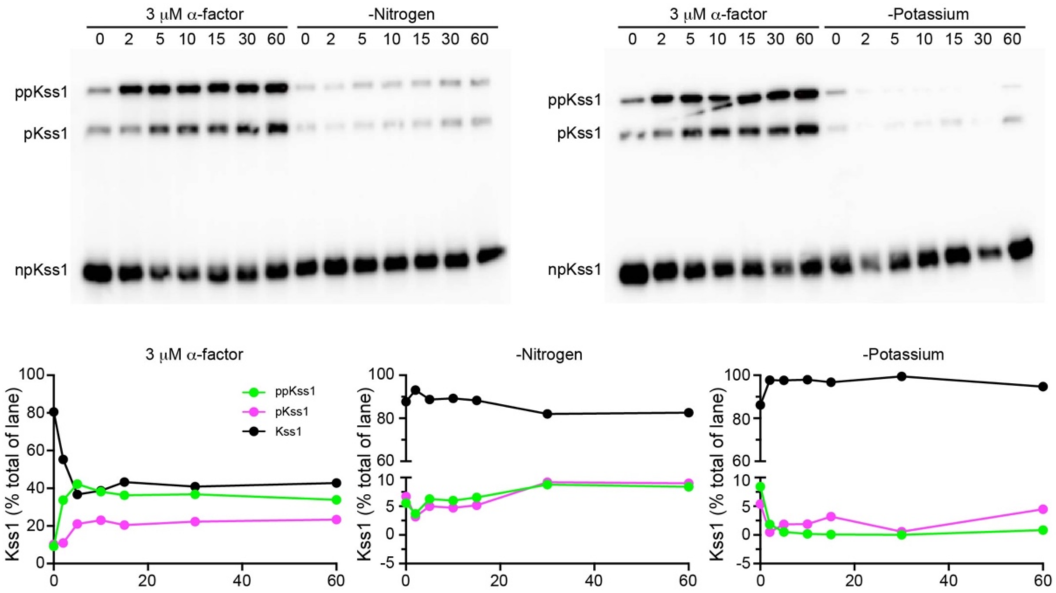


**Figure S5: Nitrogen starvation activates the MAP kinase Kss1.** Phostag immunoblotting analysis of cells starved for potassium or nitrogen, or treated with 3 μM α-factor for 60 min. Protein bands (top) representing dual (ppKss1), mono (pKss1) and nonphosphorylated (Kss1) forms of the protein were quantified using ImageLab and presented as percentage of total Kss1 at 0 min (bottom).

**FIGURE S6**

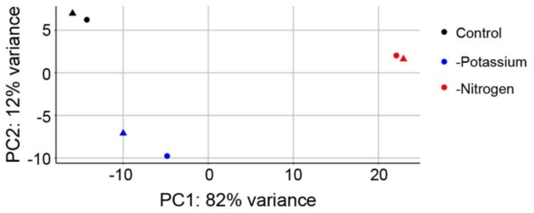


**Figure S6: Potassium and nitrogen starvation exhibit distinct transcriptional profiles.** Principal component analysis of RNA sequencing data obtained from **Figure 4A**.

**TABLE S1**

**Yeast strains used in this study**

| Strain | Description | Source |
|---|---|---|
| *BY4741* | *MAT**a** his3Δ leu2Δ met15Δ ura3Δ LYS2* | Yeast Knockout Collection (Invitrogen) |
| ***FIGURE 2*** | | |
| *pep4Δ* | *BY4741 pep4Δ::KanMX* | Yeast Knockout Collection (Invitrogen) |
| ***FIGURE 3*** | | |
| *atg1Δ* | *BY4741 atg1Δ::KanMX* | This study |
| *atg5Δ* | *BY4741 atg5Δ::KanMX* | This study |
| *vps34Δ* | *BY4741 vps34Δ::KanMX* | (1) (2) |
| *vps15Δ* | *BY4741 vps15Δ::KanMX* | |
| *vps30Δ* | *BY4741 vps30Δ::KanMX* | |
| *vps38Δ* | *BY4741 vps38Δ::KanMX* | |
| *atg14Δ* | *BY4741 atg14Δ::KanMX* | |
| *pho8*Δ60 | *BY4741 pho8Δ::pho8Δ60::KanMX* | Gift from Mara Duncan (U. Michigan) |
| *om45-GFP* | *BY4741 Om45-GFP::His3MX6* | Yeast GFP Clone Collection (Thermo Fisher Scientific) |
| *idh1-GFP* | *BY4741 Idh1-GFP::His3MX6* | Yeast GFP Clone Collection (Thermo Fisher Scientific) |
| ***FIGURE S3*** | | |
| *kss1-myc* | *BY4741 kss1-9xMyc::hphMX* | This study |

**REFERENCES**


1. Slessareva, J. E., Routt, S. M., Temple, B., Bankaitis, V. A., and Dohlman, H. G. (2006) Activation of the phosphatidylinositol 3-kinase Vps34 by a G protein alpha subunit at the endosome. *Cell* **126**, 191-203
2. Rangarajan, N., Gordy, C. L., Askew, L., Bevill, S. M., Elston, T. C., Errede, B., Hurst, J. H., Kelley, J. B., Sheetz, J. B., Suzuki, S. K., Valentin, N. H., Young, E., and Dohlman, H. G. (2019) Systematic analysis of F-box proteins reveals a new branch of the yeast mating pathway. *J Biol Chem* **294**, 14717-14731
